# The Bovine Genome Variation Database (BGVD): Integrated Web-database for Bovine Sequencing Variations and Selective Signatures

**DOI:** 10.1101/802223

**Authors:** Ningbo Chen, Weiwei Fu, Jianbang Zhao, Jiafei Shen, Qiuming Chen, Zhuqing Zheng, Hong Chen, Tad S. Sonstegard, Chuzhao Lei, Yu Jiang

**Affiliations:** Key Laboratory of Animal Genetics, Breeding and Reproduction of Shaanxi Province, College of Animal Science and Technology, Northwest A&F University, Yangling 712100, China; College of Information Engineering, Northwest A&F University, Yangling 712100, China; Recombinetics, Inc., St Paul, MN, USA

## Abstract

Next-generation sequencing has yielded a vast amount of cattle genomic data for the global characterization of population genetic diversity and the identification of regions of the genome under natural and artificial selection. However, efficient storage, querying and visualization of such large datasets remain challenging. Here, we developed a comprehensive Bovine Genome Variation Database (BGVD, http://animal.nwsuaf.edu.cn/BosVar) that provides six main functionalities: Gene Search, Variation Search, Genomic Signature Search, Genome Browser, Alignment Search Tools and the Genome Coordinate Conversion Tool. The BGVD contains information on genomic variations comprising ∼60.44 M SNPs, ∼6.86 M indels, 76,634 CNV regions and signatures of selective sweeps in 432 samples from modern cattle worldwide. Users can quickly retrieve distribution patterns of these variations for 54 cattle breeds through an interactive source of breed origin map using a given gene symbol or genomic region for any of the three versions of the bovine reference genomes (ARS-UCD1.2, UMD3.1.1, and Btau 5.0.1). Signals of selection are displayed as Manhattan plots and Genome Browser tracks. To further investigate and visualize the relationships between variants and signatures of selection, the Genome Browser integrates all variations, selection data and resources from NCBI, the UCSC Genome Browser and AnimalQTLdb. Collectively, all these features make the BGVD a useful archive for in-depth data mining and analyses of cattle biology and cattle breeding on a global scale.

## Introduction

Cattle are usually considered the most economically important livestock. The species numbers more than 1.4 billion on a global scale, constituting some 800 extant cattle breeds (FAO, 2016, http://www.fao.org/home/en/). Cattle are now kept on all inhabited continents, in contrasting climatic zones and under very different conditions [1]. The different uses of cattle and the selection for desired traits have resulted in diverse populations distributed across the world. To meet projected global demands for food, initiatives such as the cattle genome project [2–5] are generating resequencing data from breeds worldwide. The DNA-based selection tools built from these data are further accelerating rates of genetic gain and improving animal health and welfare [2]. However, the limited amount of variation data provided by dbSNP [6], restricted access to the 1000 Bull Genomes Project [7], and the existence of only sporadic cattle databases that are specialized in gene and quantitative trait locus (QTL) annotation [8–10] considerably hinder the utility of these data. Furthermore, accessing and integrating resequencing data in a highly interactive, user-friendly web interface, especially data for allele frequency resource and selection in natural populations, is a pre-requisite for identifying functional genes. Therefore, building a public data repository is vital for collecting a wide variety of cattle resequencing data and performing integrative, in-depth analyses within the research community.

Here, we develop the Bovine Genome Variation Database (BGVD), the first web-based public database for accessing dense and broadly representative bovine whole-genome variation data. The BGVD is a data repository that focuses on single nucleotide polymorphisms (SNPs), indels, copy number variations (CNVs), and selective signatures underlying domestication and population bottleneck events. We have implemented a large number of summary statistics informative for the action of selection, such as nucleotide diversity (Pi) [11], heterozygosity (*H*_p_) [12], integrated haplotype score (iHS) [13], Weir and Cockerham’s *F*_ST_ [14], cross-population extended haplotype homozygosity (XP-EHH) [15], and the cross-population composite likelihood ratio (XP-CLR) [16] (Table 1). Six early differentiated ancestral populations were used for selection analysis: African taurine, European taurine, Eurasian taurine, East Asian taurine, Chinese indicine and Indian indicine. The current version of the BGVD contains 60,439,391 SNPs, 6,859,056 indels, and 76,634 CNV regions derived from 432 cattle. With its functionalities for browsing for variations and their selection scores, the BGVD provides an important publicly accessible resource to the research community to facilitate breeding research and applications and provides information on dominant functional loci and targets for genetic improvement through selection.

**Table 1.** Statistical terms for selection sweep in the Bovine Genome Variation Database (BGVD)

| Statistical term | Abbreviation | Population 1 | Population 2 | Windows |
| --- | --- | --- | --- | --- |
| Nucleotide diversity | Pi | Indian indicine (IN) |  | 30k |
| Heterozygosity | $H_p$ | Chinese indicine (CN) | | 60k |
| Integrated haplotype score | his | East Asian taurine (EA)<br>Eurasian taurine (EUA)<br>European taurine (EUR)<br>African taurine (AFR)<br>*Bos indicus* (BIN)<br>*Bos taurus* (BTA) |  | 30k |
| Weir and Cockerham's $F_{ST}$ | $F_{ST}$ | Indian indicine (IN) | Other five groups | 30k |
| Cross-population composite likelihood ratio | XP-CLR | Chinese indicine(CN) | Other five groups | 30k |
| Cross-population extended haplotype homozygosity | XP-EHH | East Asian taurine (EA)<br>Eurasian taurine (EUA)<br>European taurine (EUR)<br>African taurine (AFR)<br>*Bos indicus* (BIN) | Other five groups<br>Other five groups<br>Other five groups<br>Other five groups<br>*Bos taurus* (BTA) | 30k |

## Database structure and content

The BGVD includes SNPs, indels, CNVs, genomic selection, and other database resources including NCBI, UCSC Genome Browser, AnimalQTLdb, KEGG, and AmiGO 2 for cattle. A detailed description is provided in the following sections and documents on the homepage.

### Sample information

Our data set integrates genomes from previously published cattle genetic works [3–5,17–21], providing a total of 432 samples representing 54 breeds. All raw sequence data were obtained from the Sequence Read Archive (SRA) of NCBI. The set of samples is grouped by location of breed origin and contains the following number of individuals: 108 West European, 83 Central-South European, 9 Middle East, 9 Tibetan, 28 Northeast Asian, 47 North-Central Chinese, 21 Northwest Chinese, 33 South Chinese, 24 Indo-Pakistan, and 70 African cattle. Geographic information and other detailed information for each sample are provided on the homepage and the corresponding ‘Sample Table’ page.

### Variants information

Data were processed and loaded into the BGVD using the following pipeline according to previously published protocols [5] (**Figure 1A**, see detailed description on the Documentation page at of the website). First, short, 250 bp paired-end Illumina reads were aligned to the Btau 5.0.1 genome assembly (GCF_000003205.7) using BWA [22], resulting in an average of ∼13X coverage of the bovine genome among the cattle varieties. Duplicate reads were removed using Picard tools (http://broadinstitute.github.io/picard/). The Genome Analysis Toolkit (GATK) was used to detect SNPs and indels [23]. A total of ∼60.4 million autosomal SNPs and ∼6.8 million autosomal indels were identified. Beagle was used to phase the identified SNPs [24]. Annotation of SNPs and indels was carried out by using snpEff [25]. Minor allele frequency (MAF) for all cattle and allele frequencies for each breed and the “core” cattle group (see Population structure section) were calculated with PLINK [26]. CNVcaller [27] was used to discover CNVs, and 76,634 CNV regions (CNVR) were detected in 432 cattle genomes. Then, the CNVs were annotated using Annovar [28]. Given that three versions of the bovine genome, Btau 5.0.1, UMD3.1.1, and the newly released ARS-UCD1.2 (project accession: NKLS00000000), are commonly used, we produced liftOver chain files (Btau5.0.1ToUMD3.1.1.chain.gz and Btau5.0.1ToARS-UCD1.2.chain.gz) and converted variation coordinates to those of the other two genomes using liftOver [29].

**Figure 1.**
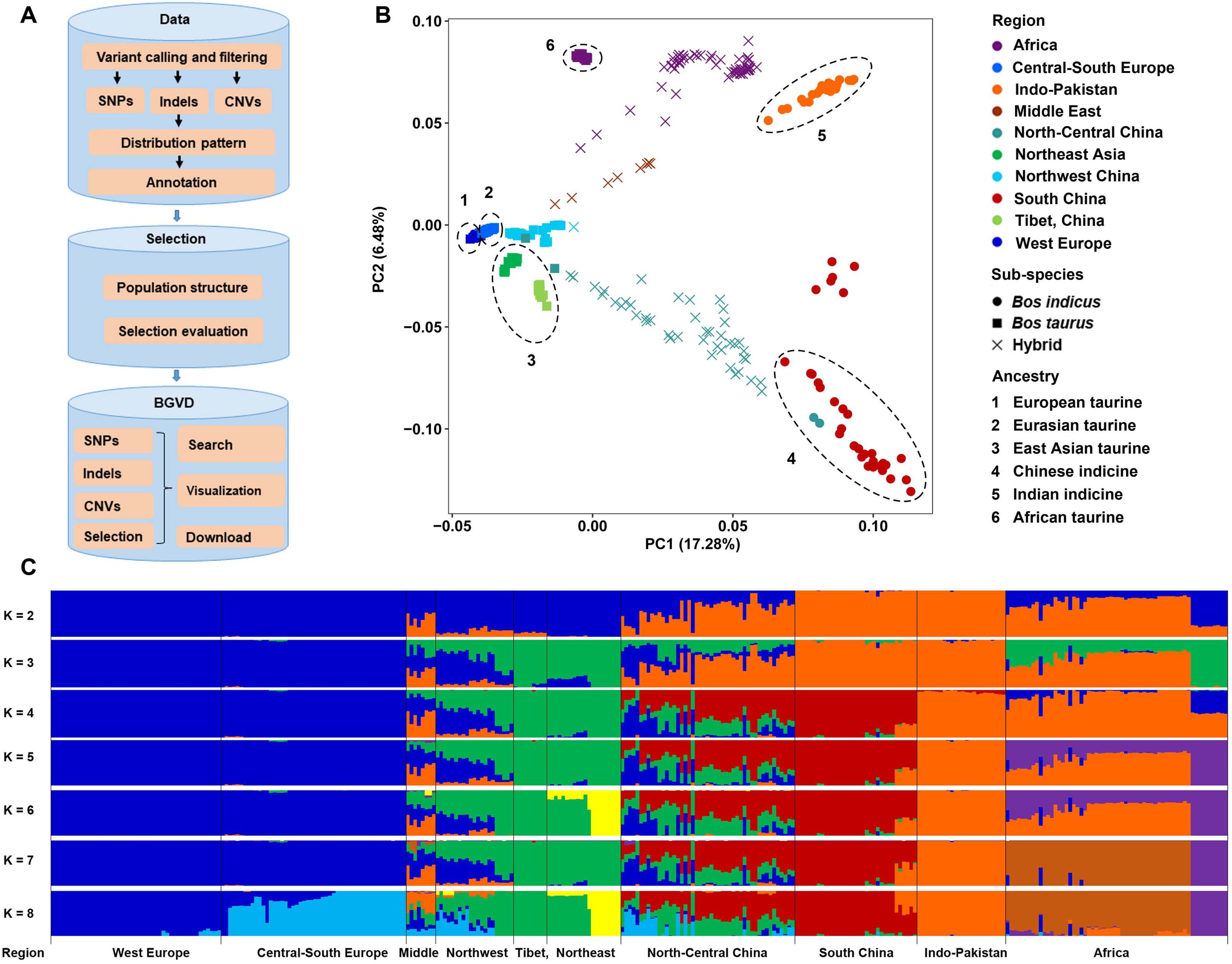
Analysis pipeline used to construct the database and population analysis of 432 cattle. **A.** Analysis pipeline used to construct the database. **B.** Principal component analysis of 432 cattle; different numbers in B represent six “core” cattle groups. **C.** Model-based clustering of cattle breeds using the program ADMIXTURE with *K* = 2 to 8 (plotted in R).

### Population structure

The population structure of all cattle was inferred using Eigensoft and ADMIXTURE [30,31], based on the genome-wide unlinked SNP dataset, all according to previously published protocols [5]. All 432 individuals were used for principal component analysis, and the results were consistent with our previous results [6], except that the African taurine cattle were split form other taurine cattle (Figure 1B). To reduce the bias due to sample size, 10 individuals were randomly selected for breeds that had more than 10 samples. A total of 317 cattle samples were selected for estimating ancestral populations by setting *K* = 2 through *K* = 8 in ADMIXTURE (Figure 1C). Combining our previous results [5], in addition to five geographically distributed ancestral groups (European taurine, Eurasian taurine, East Asian taurine, Chinese indicine, and Indian indicine), African taurine was added in this study (Figure 1B).

### Selection evaluation

The BGVD provides signatures of selection for eight groups, six of which were the “core” cattle groups that we identified as ancestral groups and the other two of which were directly divided into two categories based on sub-species: *Bos indicus* and *Bos taurus*. Here, selective signals were evaluated using six methods, namely, Pi, *H*_p_, iHS, *F*_ST_, XP-EHH, and XP-CLR (**Table 1**).

### Database implementation

The web interface of the BGVD was built by combining an Apache web server, the PHP language, HTML, JavaScript, and the relational database managements system MySQL. High-quality SNPs, indels, CNVs, selection scores and their corresponding annotations, classification and threshold values, were processed with Perl scripts and stored in the MySQL database. The server application was written in PHP, and CodeIgniter was chosen as the model-view-controller (MVC) framework for the system. A client interface developed with HTML5 and JavaScript was used to implement search, data visualization and download. Moreover, we introduced web-based software such as BLAST, BLAT, liftOver, and the UCSC Genome Browser (hereafter referred to as ‘Gbrowse’) [29,32] into the BGVD. Information including variations, selection scores, gene annotation, QTLs, and phastCons conserved elements of 20-way mammals and 100-way vertebrates was integrated into Gbrowse to facilitate global presentation.

## Web interface and usage

The BGVD uses a series of user-friendly interfaces to display results. All the parts in our browser are dynamic and interactive. We provided six main functionalities: (i) Gene Quick Search, (ii) Variation Search, (iii) Genomic Selection Search, (iv) Genome Browser, (v) Alignment Search Tools (BLAT/BLAST), and (vi) Genome Coordinate Conversion Tool (liftOver).

For “Gene Quick Search”, we integrated information from NCBI, AmiGO 2, and KEGG. Users can input a gene symbol to view all available information, including basic gene information (*e.g.*, genomic location, transcript and protein profile, relevant Gene Ontology (GO) ID, GO terms, and KEGG pathways), gene variations (*e.g.*, SNPs, indels, and CNVs), as well as selective signatures. We also provide links to Gbrowse and external databases (NCBI, AmiGO 2, and KEGG) to help the user obtain more information, such as gene/mRNA/protein sequence, KEGG Orthology (KO), and motif.

For “Variation Search”, the BGVD allows users to obtain information on SNPs, indels, and CNVs by searching for a specific gene or a genomic region in three versions of the bovine genome (ARS-UCD1.2, UMD3.1.1, and Btau 5.0.1) (**Figure 2A**). Users can filter SNPs and indels further by “Advanced Search”, in which certain parameters (Figure 2B), such as MAF and consequence type, can be set; this option enables users to narrow down the items of interest in an efficient and intuitive manner.

**Figure 2.**
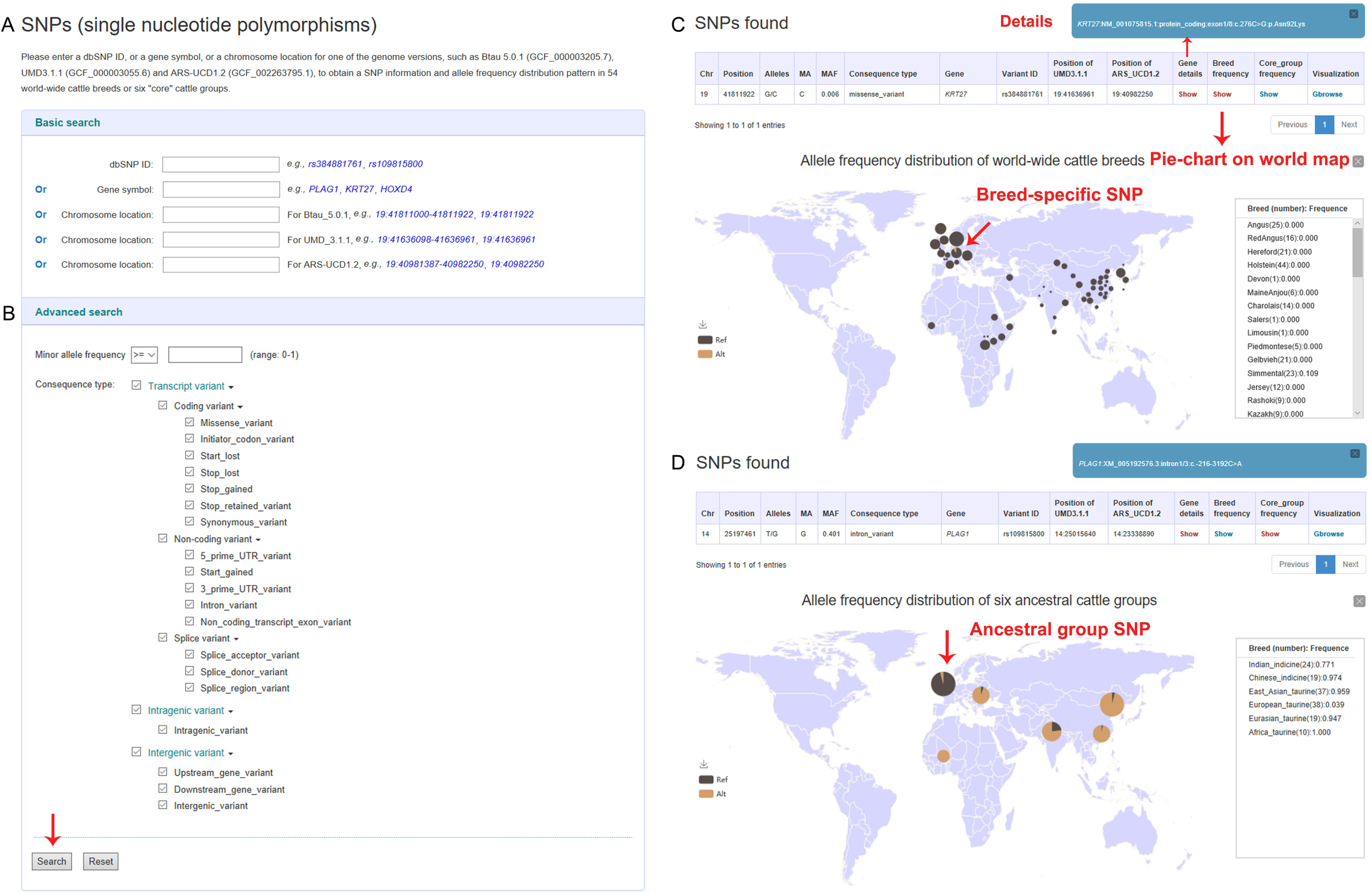
Screenshots of a single nucleotide polymorphism (SNP) data search and the results for two examples. **A.** Search items involving rs ID, gene name and position of three bovine reference genomes. **B.** Advanced Search menu enabling filtering for minor allele frequency and consequence type. **C.** Detailed annotation of the rs384881761 locus of the *KRT27* gene and the allele frequency distribution pie-chart of 54 cattle breeds worldwide. **D.** Display format of the allele frequency for the rs109815800 locus of *PLAG1* among defined ancestral groups.

The results are presented in an interactive table and graph. For SNPs and indels, users can obtain related details including variant position, alleles, MAF, variant effect, rs ID and the allele frequency distribution pattern in 54 cattle breeds worldwide (Figure 2C) or in six “core” cattle groups (Figure 2D), which could help users dynamically visualize breed-specific (rs384881761, *KRT27*) [2] or ancestral group-specific (rs109815800, *PLAG1*) [33] variants and their global geographical distributions.

For CNVs, users can obtain information about CNVR, such as intersected genomic regions, CNV length, the closest gene, consequence type (**Figure 3A**), and copy number distribution in 432 individuals representing 49 cattle populations. We provide three types of display formats of copy number distributions in which the categories and haploid copy number of each individual can be viewed (Figure 3B-D), such as the “view” button, which produces a scatterplot (*MATN3*); “Gbrowse”, which is linked to the “CNVR Bar” track (*KIT*); and the more detailed visualization “cnvBar” track, which generates a box-whisker plot (*CIITA*) [34].

**Figure 3.**
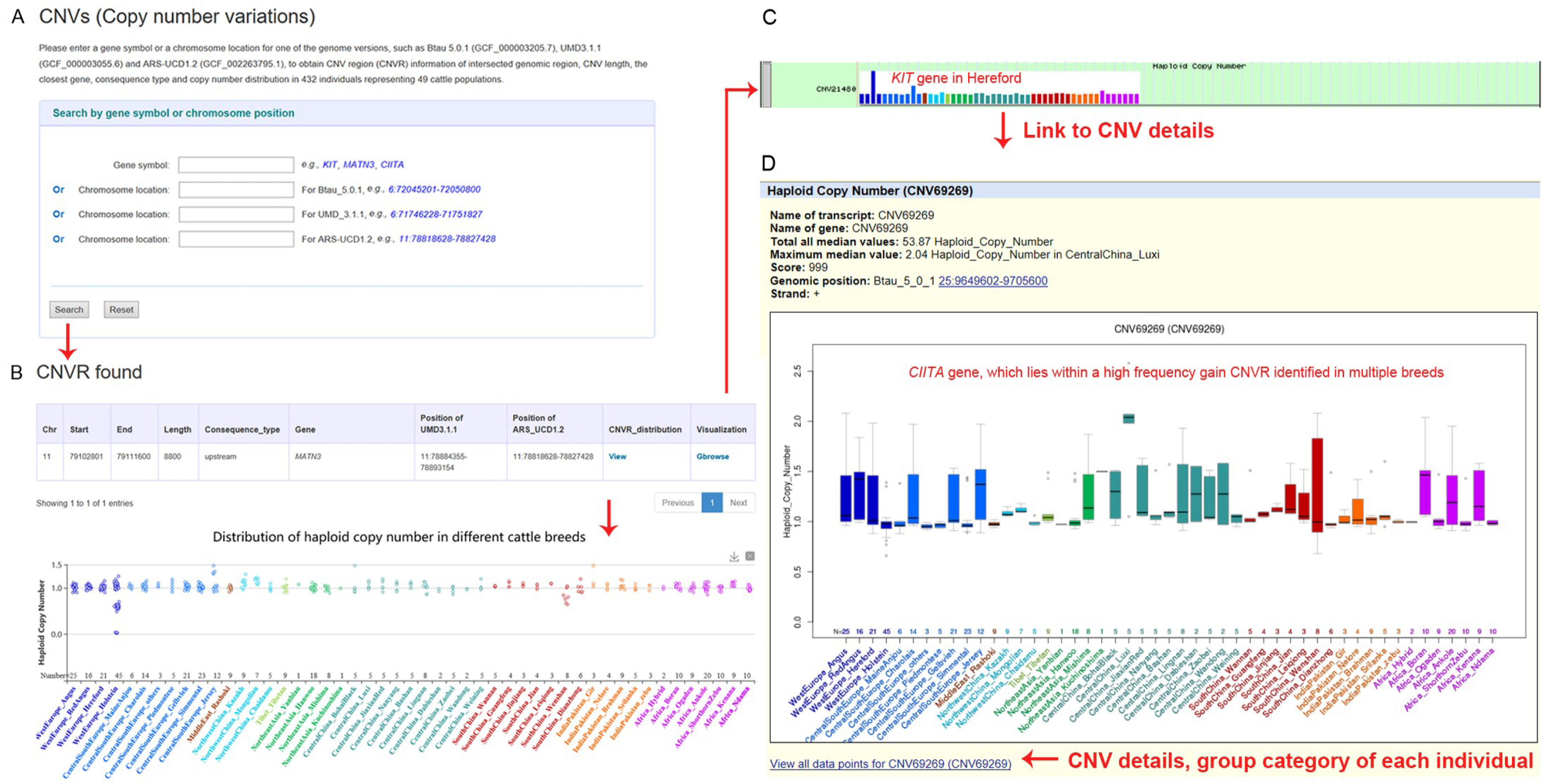
Screenshots of a copy number variation region (CNVR) data search and three types of display formats of the results. **A.** Search items involving the gene name and position of three bovine reference genomes. **B.** Results involving detailed annotation for the CNVR and copy number distribution patterns of 432 individuals representing 49 populations. An example of *MATN3*, which showed different copy numbers in the Holstein population. **C.** “CNVR Bar” track in the bar chart format in UCSC Genome Browser (Gbrowse). An example of the *KIT* gene, which is related to coat color in Herefords. **D.** The more detailed visualization “CNVR Bar” track in the format of a box-whisker plot, displaying copy number distribution in 49 cattle populations. An example of *CIITA*, which lies within a high-frequency gain CNVR identified in multiple breeds that showed nematode resistance.

In the genomic signature interface, users can select a specific gene symbol or genomic region, one of the statistical methods (Pi, *H*_p_, iHs, *F*_ST_, XP-CLR, or XP-EHH), and a specific “core” cattle group to view the selection scores (Table 1 and **Figure 4A**). In our database, the selection scores are pre-processed by several algorithms (Z-transform and logarithm). The results are retrieved in a tabular format (Figure 4B). When users click the “show” button on the table, selective signals are displayed in Manhattan plots or common graphics, where the target region or gene is highlighted in a red/blue colour. In addition, the “Gbrowse” button can locate the position of the selection and differentiation profiles of specific groups (Figure 4C). To demonstrate the function of our database, we extracted results for a number of putatively selected genes detected by different methods: *OR2T33* [35] (Figure 4B, C), *STOM, EPB42* [3], *PLAG1* [33], *MSRB3* [35], *CDC42SE1* [36], *R3HDM1* [37], and *ASIP* [5] (Figure 4C).

**Figure 4.**
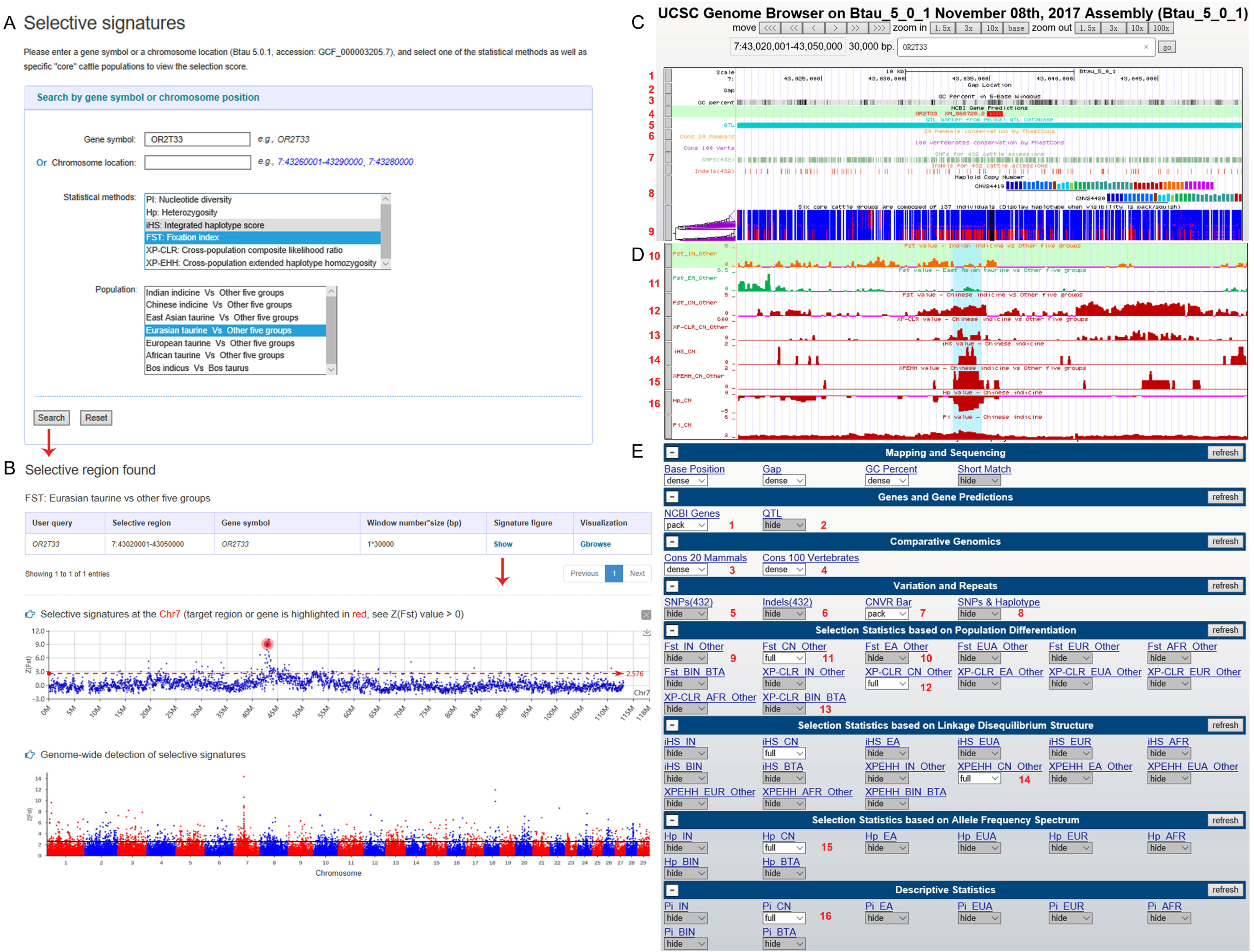
Screenshots of a search for genomic selection data and representation of the selection data. **A.** Search items involving gene name, position, and one of the statistical methods (nucleotide diversity (Pi), heterozygosity (Hp), integrated haplotype score (iHS), Weir and Cockerham’s *F*_ST_, cross-population extended haplotype homozygosity (XP-EHH), and the cross-population composite likelihood ratio (XP-CLR)), and specific “core” cattle groups. **B.** Detailed annotation for the target gene or region in the variant grid and the corresponding selective signal at the chromosome and whole-genome levels, respectively. An example of selective signal of the *OR2T33* gene in Eurasian taurine population. **C.-E.** The display of 57 tracks in UCSC Genome Browser (Gbrowse) in the BGVD. Numbers 1-16 represent the corresponding tracks. (C) Example of the *OR2T33* gene in “SNPs&Hap” track. Different haplotypes of the *Bos taurus* and *Bos indicus* groups are shown in blue and red, respectively. D. Examples of the six selection scores of the *POFUT1* gene in the Chinese indicine (CN) group, and where each group is represented by a different color. Here, we show *F*_ST_ scores of Indian indicine (IN) and East Asian (EA) groups with orange and blue, respectively. E. Fifty-seven tracks in Gbrowse.

To further investigate the relationship between variations and signatures of selection, Gbrowse has been introduced to support our database. Currently, 57 tracks have been released for the Btau 5.0.1 assembly. Users can search with a gene symbol or genomic region to see SNPs, indels, CNVs, genomic signatures, QTLs, and conserved elements in the global view. All search pages in the BGVD allow quick access to Gbrowse to deepen the functional inference of the candidate gene or region by combining other tracks. Most noteworthy, the phased haplotypes from six “core” cattle groups are displayed in “SNPs&Hap” track. The ‘squish’ or ‘pack’ view highlights local patterns of genetic linkage between variants. In the haplotype sorting display, variants are presented as vertical bars with reference alleles in blue and alternate alleles in red so that local patterns of linkage can be easily discerned when clustering is used to visually group co-occurring allele sequences in haplotypes. We display different haplotypes of the *Bos taurus* and *Bos indicus* groups in Figure 4C. We highlight that the tracks of selection statistics from different populations are visualized in different colours (Figure 4D).

We also introduced two sequence alignment tools, webBlat, and NCBI wwwBLAST, as well as a genome coordinate conversion tool (liftOver) [29] into the BGVD. The webBlat tool can be used to quickly search for homologous regions of a DNA or mRNA sequence, which can then be displayed in Gbrowse. BLAST can find regions of local similarity between sequences, which can be used to infer functional and evolutionary relationships between sequences. The liftOver tool is used to translate genomic coordinates from one assembly version into another. Our database provides an online lift from Btau_5.0.1 to UMD_3.1.1 and from Btau_5.0.1 to ARS-UCD1.2.

## Discussion

By applying summary statistics to a relatively extensive data set from cattle genomes, we provide a timely and expandable resource for the population genomics research community. An associated user-friendly genome browser gives a representation of the genetic variation in a genomic region of interest and offers functionality for an array of downstream analyses. We expect that the database will prove useful for genome mining through the large number of test statistics and the fine-grained character of resequencing data. We believe that this expandable resource will facilitate the interpretation of signals of selection at different temporal, geographical and genomic scales.

## Authors’ contributions

NC, WF, and YJ conceived of the project and designed the research. NC and WF drafted the manuscript. TS, CL, YJ, HC, and ZZ revised the manuscript. NC, JS, and QC performed the data analyses. WF and JZ wrote the source code for the BGVD.

## Competing interests

The authors declare that they have no competing interests.

## Acknowledgments

The project was supported by the National Natural Science Foundation of China (31822052), the National Thousand Youth Talents Plan (both to Yu Jiang), the National Beef Cattle and Yak Industrial Technology System (Grant No. CARS-37) and the National Natural Science Foundation of China (Grant No. 31872317) (both to Chuzhao Lei).

## Notes

http://animal.nwsuaf.edu.cn/code/index.php/BosVar

## References

[1] Felius M, Koolmees PA, Theunissen B, Lenstra JA. On the breeds of cattle—historic and current classifications. Diversity 2011;3:660–92.

[2] Daetwyler HD, Capitan A, Pausch H, Stothard P, Van Binsbergen R, Brondum RF, et al. Whole-genome sequencing of 234 bulls facilitates mapping of monogenic and complex traits in cattle. Nat Genet 2014;46:858–65.

[3] Kim J, Hanotte O, Mwai O, Dessie T, Bashir S, Diallo B, et al. The genome landscape of indigenous African cattle. Genome Biol 2017;18:34.

[4] Stothard P, Liao X, Arantes AS, De Pauw M, Coros C, Plastow G, et al. A large and diverse collection of bovine genome sequences from the Canadian Cattle Genome Project. GigaScience 2015;4:49.

[5] Chen N, Cai Y, Chen Q, Li R, Wang K, Huang Y, et al. Whole-genome resequencing reveals world-wide ancestry and adaptive introgression events of domesticated cattle in East Asia. Nat Commun 2018;9:2337.

[6] Cunningham F, Achuthan P, Akanni W, Allen J, Amode M R, Armean IM, et al. Ensembl 2019. Nucleic Acids Res 2018;47:746–51.

[7] Hayes BJ, MacLeod IM, Daetwyler HD, Phil BJ, Chamberlain AJ, Vander Jagt C, et al. Genomic prediction from whole genome sequence in livestock: the 1000 bull genomes project. 10th World Cong Genet Appl Livestock Produc (WCGALP) 2014.

[8] Song S, Tian D, Li C, Tang B, Dong L, Xiao J, et al. Genome Variation Map: a data repository of genome variations in BIG Data Center. Nucleic Acids Res 2018;46:944–9.

[9] Elsik CG, Unni D, Diesh C, Tayal A, Emery ML, Nguyen HN, et al. Bovine Genome Database: new tools for gleaning function from the Bos taurus genome. Nucleic Acids Res 2016;44:834–9.

[10] Childers CP, Reese JT, Sundaram JP, Vile DC, Dickens CM, Childs KL, et al. Bovine Genome Database: integrated tools for genome annotation and discovery. Nucleic Acids Res 2011;39:830–4.

[11] Nei M, Li W. Mathematical model for studying genetic variation in terms of restriction endonucleases. Proc Natl Acad Sci U S A 1979;76:5269–73.

[12] Rubin C, Zody MC, Eriksson J, Meadows JRS, Sherwood E, Webster MT, et al. Whole-genome resequencing reveals loci under selection during chicken domestication. Nature 2010;464:587–91.

[13] Voight BF, Kudaravalli S, Wen X, Pritchard JK. A map of recent positive selection in the human genome. PLoS Biol 2006;4:446–58.

[14] 28. Weir BS, Cockerham CC. Estimating F-statistics for the analysis of populaition structure. Evolution 1984;38:1358–70.

[15] Sabeti PC, Varilly P, Fry B, Lohmueller J, Hostetter E, Cotsapas C, et al. Genome-wide detection and characterization of positive selection in human populations. Nature 2007;449:913–8.

[16] Chen H, Patterson N, Reich D. Population differentiation as a test for selective sweeps. Genome Res 2010;20:393–402.

[17] Heaton MP, Smith TPL, Carnahan JK, Basnayake V, Qiu J, Simpson B, et al. Using diverse U.S. beef cattle genomes to identify missense mutations in EPAS1, a gene associated with pulmonary hypertension. F1000Research 2016;5:2003.

[18] Bickhart DM, Xu L, Hutchison JL, Cole JB, Null DJ, Schroeder SG, et al. Diversity and population-genetic properties of copy number variations and multicopy genes in cattle. DNA Res 2016;23:253–62.

[19] Shin D, Lee HJ, Cho S, Kim HJ, Hwang JY, Lee C, et al. Deleted copy number variation of Hanwoo and Holstein using next generation sequencing at the population level. BMC genomics 2014;15:240.

[20] Tsuda K, Kawaharamiki R, Sano S, Imai M, Noguchi T, Inayoshi Y, et al. Abundant sequence divergence in the native Japanese cattle Mishima-Ushi (Bos taurus) detected using whole-genome sequencing. Genomics 2013;102:372–8.

[21] Kawaharamiki R, Tsuda K, Shiwa Y, Araikichise Y, Matsumoto T, Kanesaki Y, et al. Whole-genome resequencing shows numerous genes with nonsynonymous SNPs in the Japanese native cattle Kuchinoshima-Ushi. BMC genomics 2011;12:103–10.

[22] Li H. Exploring single-sample SNP and INDEL calling with whole-genome de novo assembly. Bioinformatics 2012;28:1838–44.

[23] Mckenna A, Hanna M, Banks E, Sivachenko A, Cibulskis K, Kernytsky AM, et al. The Genome Analysis Toolkit: a MapReduce framework for analyzing next-generation DNA sequencing data. Genome Res 2010;20:1297–303.

[24] Browning SR, Browning BL. Rapid and accurate haplotype phasing and missing-data inference for whole-genome association studies by use of localized haplotype clustering. Am J Hum Genet 2007;81:1084–97.

[25] Cingolani P, Platts AE, Wang LL, Coon M, Nguyen T, Wang L, et al. A program for annotating and predicting the effects of single nucleotide polymorphisms, SnpEff: SNPs in the genome of Drosophila melanogaster strain w1118; iso-2; iso-3. Fly 2012;6:80–92.

[26] Purcell S, Neale BM, Toddbrown K, Thomas L, Ferreira MAR, Bender D, et al. PLINK: a tool set for whole-genome association and population-based linkage analyses. Am J Hum Genet 2007;81:559–75.

[27] Wang X, Zheng Z, Cai Y, Chen T, Li C, Fu W, et al. CNVcaller: highly efficient and widely applicable software for detecting copy number variations in large populations. GigaScience 2017;6:1–12.

[28] Wang K, Li M, Hakonarson H. ANNOVAR: functional annotation of genetic variants from high-throughput sequencing data. Nucleic Acids Res 2010;38:e164.

[29] Casper J, Zweig AS, Villarreal C, Tyner C, Speir ML, Rosenbloom KR, et al. The UCSC Genome Browser database: 2018 update. Nucleic Acids Res 2018;46:762–9.

[30] Patterson N, Price AL, Reich D. Population structure and eigenanalysis. PLoS Genet 2006;2:e190.

[31] Alexander DH, Novembre J, Lange K. Fast model-based estimation of ancestry in unrelated individuals. Genome Res 2009;19:1655–64.

[32] Geer LY, Marchlerbauer A, Geer RC, Han L, He J, He S, et al. The NCBI BioSystems database. Nucleic Acids Res 2010;38:492–6.

[33] Bouwman AC, Daetwyler HD, Chamberlain AJ, Ponce CH, Sargolzaei M, Schenkel FS, et al. Meta-analysis of genome-wide association studies for cattle stature identifies common genes that regulate body size in mammals. Nat Genet 2018;50:362–7.

[34] 48. Liu GE, Brown T, Hebert DA, Cardone MF, Hou Y, Choudhary RK, et al. Initial analysis of copy number variations in cattle selected for resistance or susceptibility to intestinal nematodes. Mamm Genome 2011;22:111–21.

[35] Ramey HR, Decker JE, Mckay SD, Rolf MM, Schnabel RD, Taylor JF. Detection of selective sweeps in cattle using genome-wide SNP data. BMC genomics 2013;14:382.

[36] Portoneto LR, Sonstegard TS, Liu GE, Bickhart DM, Silva MVGBD, Machado MA, et al. Genomic divergence of zebu and taurine cattle identified through high-density SNP genotyping. BMC genomics 2013;14:876.

[37] Gibbs R, Taylor J, Van Tassel C, Barendse W, Eversole K, Gill C, et al. Genome-wide survey of SNP variation uncovers the genetic structure of cattle breeds. Science 2009;324:528–32.

